# Fibroblast Growth Factor 2 lethally sensitizes cancer cells to stress-targeted therapeutic inhibitors

**DOI:** 10.1101/227496

**Authors:** Matheus H. Dias, Cecília S Fonseca, Julianna D. Zeidler, Layra L. Albuquerque, Marcelo S. da Silva, Eduardo Cararo-Lopes, Marcelo S. Reis, Vincent Noël, Ian A. Prior, Hugo A. Armelin

## Abstract

In malignant transformation, cellular stress response pathways are dynamically mobilized to counterbalance oncogenic activity, keeping cancer cells viable. Therapeutic disruption of this riskily balanced homeostasis might change the outcome of many human cancers, particularly those for which no effective therapy is available. Here, we report the use of Fibroblast Growth Factor 2 (FGF2) to demonstrate that further mitogenic activation disrupts cellular homeostasis and strongly sensitizes cancer cells to stress-targeted therapeutic inhibitors. We show that FGF2 enhanced replication and proteotoxic stresses in a K-Ras-driven murine cancer cell model, and combinations of FGF2 and proteasome or DNA damage response-checkpoint inhibitors triggered cell death. CRISPR/Cas9-mediated K-Ras depletion suppressed the malignant phenotype and prevented these synergic toxicities in these murine cells. Moreover, in a panel of human Ewing’s sarcoma family tumor cells, sub-lethal concentrations of bortezomib (proteasome-inhibitor) or VE-821 (ATR-inhibitor) induced cell death when combined with FGF2. Sustained MAPK-ERK1/2 overactivation induced by FGF2 underlies these synthetic lethalities, once late pharmacological inhibition of this pathway restored cell homeostasis and prevented these described synergies. Our results highlight how mitotic signaling pathways frequently overridden in malignant transformation might be exploited to disrupt the risky robustness of cancer cells, ultimately sensitizing them to stress-targeted therapies. This approach provides a new therapeutic rationale for human cancers, with important implications for tumors still lacking effective treatment, and for those that frequently relapse after available therapies.

## 1. INTRODUCTION

Several cellular stress response pathways are frequently mobilized in malignant cells to cope with an aggressive and highly proliferative phenotype. Identification and targeting of cancer cells-specific vulnerabilities resulting from those stresses is a promising therapeutic approach; particularly for cancers in which the driver oncogene is not clinically druggable. For instance, gain-of-function mutations or overexpression of RAS family members (KRAS, HRAS, and NRAS) are among the most prevalent oncogenic lesions in human cancers (Prior et al., 2012); and high levels of Ras activity are necessary to maintain the transformed phenotype in some Ras-driven cancers (Singh et al., 2009). Similar oncogene addiction is also described for Ewing’s Sarcoma Family Tumors (ESFT), which are a group of childhood and adolescence poorly differentiated cancers, arising from bone and soft tissues (Biswas and Bakhshi, 2016). The (11;22) (q24;q12) chromosomal translocation encoding the fused transcription factor EWS-FLI-1 is present in approximately 85% of all Ewing’s Sarcoma Family Tumor specimens and is established as the driver oncogenic lesion in these tumors (Toomey et al., 2010). In common with Ras-driven tumors, Ewing’s sarcoma tumors display a poor prognosis at metastatic stage (cure rate of 20-40%) and the lack of clinically effective targeted therapies (Gaspar et al., 2015). Hence, selective targeting of stress-response pathways supporting Ras and EWS-FLI-1-driven tumorigenesis might be game-changing for the therapy of these aggressive malignancies.

Enhanced DNA damage and replication stress are probably the best characterized and exploited stresses resulting from malignant transformation induced by Ras, EWS-FLI-1 and other oncogenes (Hills & Diffley, 2014). Genotoxic agents such as ionizing radiation (IR), cisplatin and gemcitabine are widely used in cancer therapy aiming to push tumor DNA damage/replication stress over a lethal threshold (Swift & Golsteyn, 2014). More recently, checkpoint inhibition was shown to increase the cell death induced by these genotoxic agents (Prevo et al., 2012).

The enhanced proteotoxic stress frequently found in malignant cells is also a clinical target in cancer therapy. Proteasome inhibition resulting in lethal proteotoxic stress is a protagonist treatment for some hematological cancers (Csizmar et al., 2016). Combined induction of protein misfolding further enhanced proteotoxic stress, increasing the cytotoxicity of proteasome inhibition in vitro and in vivo (Neznanov et al., 2011). Drawbacks of these stress-targeted therapies include the high overall toxicity and acquired resistance of genotoxic agents and proteasome inhibitors like bortezomib, limiting the therapeutic window and efficacy of these approaches (Kalal et al., 2017; Cavaletti & Jakubowiak, 2010). Altogether, these observations point that overload of replication or proteotoxic stress, especially in combination with the respective sensitizing inhibition, might efficiently target cancer cells-specific vulnerabilities. Therefore, identification of effective combinations “Stress induction/sensitizing inhibition” targeting selectively malignant cells is paramount.

In this regard, exogenous administration of the Fibroblast Growth Factor 2 (FGF2) might be a viable alternative to overload cellular stress pathways in cancer cells. FGF2 is the seminal member of a large family of signaling factors of undisputed importance for neurogenesis, morphogenesis, wound healing and angiogenesis, among other functions (Armelin, 1973; Itoh & Ornitz, 2011). Despite the many different pro-tumor roles attributed to FGF2 signaling (reviewed by Turner & Grose, 2010), a set of articles unequivocally demonstrate that FGF2 can also induce cytostatic and cytotoxic responses in different cancer cells, both in vivo and in vitro (Wang et al., 1998; Sturla et al., 2000; Williamson et al., 2004; Fogarty et al., 2007). In this last context, we have also previously shown that FGF2 restrains the proliferation of murine malignant cells, in which wild-type Kras is highly amplified and overexpressed (Costa et al., 2008; Salotti et al., 2013). Because FGF2 is an activator of mitogenic signaling pathways, we hypothesized that the toxicity induced by this growth factor in cancer cells might also intensify the mobilization of stress pathways, further increasing their dependency on these pathways for cell viability.

Here, we tested whether FGF2 can selectively sensitize cancer cells to stress targeted therapeutic inhibitors. We found that in K-Ras-driven mouse Y1 malignant cells FGF2 stimulation disrupts proteostasis and enhances tonic replication stress and DNA damage response (DDR) activation. Concomitant proteasome or checkpoint inhibition induced cell death in a K-Ras-dependent manner. Importantly, in human ESFT cells, combined FGF2 signaling activation and sub-lethal doses of proteasome or checkpoint inhibitors also triggered cell death. Moreover, FGF2 induced sustained MAPK-ERK1/2 overactivation in Y1 and ESFT cells; and K-Ras depletion or late pharmacological MAPK inhibition, respectively, prevented FGF2 sensitization to these stress-targeted inhibitors. These findings indicate that further mitogenic signaling activation can be employed to overload selectively cancer cells stress response pathways, disrupting their homeostatic robustness, and increasing the cytotoxicity of stress-targeted therapies.

## 2 MATERIALS AND METHODS

### 2.1 Cell lines, cell culture and treatments

The Y1 murine adrenocortical carcinoma cell line was obtained from ATCC. Y1 cells were grown at 37 °C in a 5% CO2 atmosphere in Dulbecco’s modified Eagle’s medium DMEM (Gibco) supplemented with 10% fetal bovine serum (FBS). The Y1D1 subline (Schwindt et al, 2003) was cultured in the same conditions as Y1, and the growth medium was supplemented with 0.1 mg/mL geneticin (G418; Invitrogen). Whenever G0/G1 synchronization by serum starvation was necessary DMEM-FCS medium was removed, plates were washed with PBS, and cells were grown in FCS-free DMEM for 48h prior any stimulation. ESFT cells (A673, RD-ES, SK-N-MC and TC-32) were kindly given by Professor Susan Burchill. A673 and SK-N-MC cells were grown in DMEM 10% FBS, TC-32 cells were grown in RPMI 1640 10% FBS and RD-ES cells were grown in RPMI 1640 15% FBS. Where indicated, cells were treated with recombinant human FGF2 protein (Abcam ab9596), Colchicine (Sigma C9754); Bortezomib (Selleckchem #S1013); ATM inhibitor KU55933 (Selleckchem #S1092), ATR inhibitor VE-821 (Selleckchem #S8007) and MEK inhibitor Selumetinib (AZD6244) (Selleckchem #S1008).

### 2.2 Cell cycle analysis

For BrdU/Propidium Iodide cell cycle analyses, after the indicated treatments, cells were resuspended and fixed in ice-cold 75% ethanol in PBS overnight at 4 °C. BrdU was added at 50 μM for 30 minutes before harvesting. Fixed cells were treated with 2 M HCl and 0.5% Tween-20 for 15 min for DNA denaturation and then washed sequentially with 0.1 M sodium tetraborate (pH 9.5) and ice-cold PBS. Cells were incubated with Alexa Fluor^®^ 488 anti-BrdU (Invitrogen B35130) and subsequently treated with 10 mg/mL RNase A and stained with 50 mg/mL propidium iodide in PBS for 20 minutes before analysis in the flow cytometer.

For all flow cytometer experiments, data were acquired with Attune NxT flow cytometer (Life Technologies) and analyzed with FlowJo V.10 software (Treestar, INC.). At least 20000 cells per sample were analyzed.

### 2.3 Histones on cell cycle assays

For phospho-histone H3 (S10) or phospho-H2AX-S139 (γ-H2AX) stains along cell cycle, after treatment cells were fixed as described above, washed in PBS and incubated for 1h with the conjugated histone antibodies (histone H3 S10 Millipore 06-570-AF488 or γ-H2AX Thermo Fisher 53-9865-82) Samples were then treated with 10 mg/ml RNase A and stained with 50 mg/ml propidium iodide in PBS for 20 minutes prior to analysis by flow cytometry.

### 2.4 Protein per cell assay

To measure protein/cell, 2,5×10^4^ cells per 35 mm dish were plated, let to adhere overnight and then serum starved as described for synchronization. After 48h, cells were stimulated with FCS in presence or absence of FGF2 as indicated for each experiment. For each condition and time point we harvested three plates for counting cells, as described below for growth curves, and three plates for measuring total protein concentration. To estimate the amount of protein per cell we measured the amount of protein from each plate using Bradford method and divided by the number of cells counted from each plate of the same time point and condition.

### 2.5 Western blots

Antibodies for Western blot Western blot were as follows: IRE1α (3294 Cell signaling), Bip (3183 Cell signaling), phospho-S6 Ser235/236 (4856 Cell Signaling), phospho-eIF4E Ser209 (9741 Cell Signaling), α-Tubulin (sc-8035 Santa Cruz) phospho-H2AX S139 (ab11174 Abcam), ChK1 (ab47574 Abcam), phospho-ChK1 S345 (sc-17922 Santa Cruz), phospho-ChK2 T383 (ab59408 Abcam), p38 (9212 Cell Signaling); phospho-p38 T180/Y182 (sc-15852-R Santa Cruz), p21 (sc-397 Santa Cruz), HPRT (sc-20975 Santa Cruz), K-Ras (sc-30 Santa Cruz), Actin (ab6276 Abcam), phospho-ERK Thr202/204 (4370 Cell Signaling) and ERK (4695 Cell Signaling). Analysis was performed by standard methods using enhanced chemiluminescence or fluorescence. Images were obtained using Uvitec Alliance 9.7 documentation system (Uvitec) or Odyssey system (Licor) according to the manufacturer’s settings.

### 2.6 Cell death assay

For AnnexinV/PI stain cells were plated on 35 mm dishes and treated as described for each experiment. Then, culture media were collected and reserved, plates were washed with 450 μl of PBS, which was also reserved on the respective tubes and cells were released with 150 μl of trypsin for up to 5 minutes. Cells were then suspended and homogenized with the respective culture medium/PBS. The volume of all the suspensions were adjusted to 2 ml. 250 ul of each cell suspension were mixed 1:1 with 2x Annexin binding buffer (300 mM NaCl, 2 mM MgCl_2_, 10 mM KCl, 3,6 mM CaCl_2_, 20 mM HEPES pH 7,4) containing 1:10000 AnnexinV-FITC (produced and kindly given by Dr. Shankar Varadarajan’s lab). After 10 minutes, 5 μl of PI 50 μg/ml were added to each tube and mixed by inversion. Fixed volumes of these cell suspensions were then analyzed using Attune NxT Flow cytometer (Thermo Fisher Scientific Invitrogen) allowing combined determination of cells viability and total amount of cells per plate whenever necessary. At least 20000 cells were analyzed for each individual sample.

### 2.7 Detection of BrdU foci under native DNA conditions

For detection of long fragments of single stranded DNA (ssDNA), characteristics of replication stress, exponentially growing cells in coverslips incorporated bromodeoxyuridine (BrdU) at 50 mM for 48 hours into DNA. After that, we washed the coverslips and added fresh DMEM medium with or without 10 ng/mL FGF2 for 24 hours. To ensure that all cells incorporated BrdU, 1 additional coverslip for each condition analyzed were prepared to be subjected to DNA denaturation using 2 M HCl. Next, cells were fixed using 4% of paraformaldehyde in PBS and permeabilized with 0.2% Triton-X 100. BrdU was detected (when accessible) using α-BrdU-rat (ab6326 Abcam) followed by secondary antibody goat anti-rat conjugated to Alexa Fluor 488 (A-11006 Thermo Scientific). Stained coverslips were mounted with VECTASHIELD^®^ Mounting Medium with DAPI (Vector Labs). Images were captured using Olympus BX51 fluorescence microscope coupled with a digital camera (XM10, Olympus), and analyzed using Olympus Cell F software (version 5.1.2640). At least 65 cells were analyzed per coverslip.

### 2.8 Cas9-mediated K-Ras depletion

Because Y1 cells carry genomic amplification of k-ras gene (tens of copies), we designed three specific gRNAs against k-ras using CRISPR design tool (http://crispr.mit.edu/). A scramble sequence was also designed for control. Sequences are shown below.

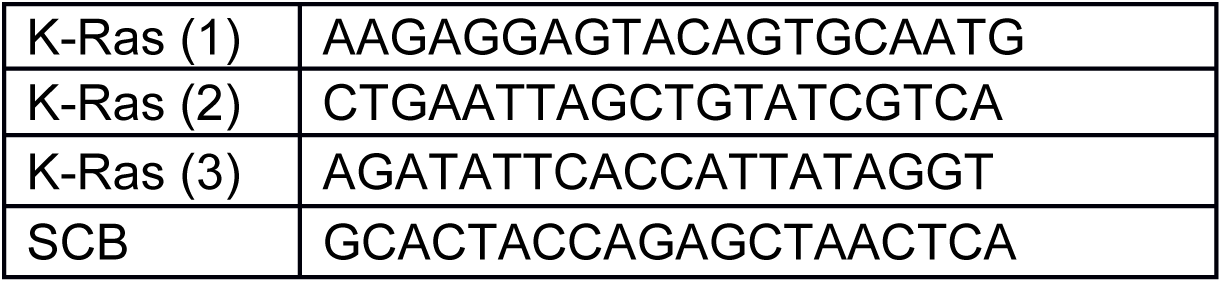

Oligos were cloned in LentiCRISPR v2 plasmid (a gift from Feng Zhang, Addgene plasmid # 52961) according to described by Sanjana and co-workers (Sanjana et al., 2014). For lentivirus production, LentiCRISPR v2 constructs, psPAX2 (a gift from Didier Trono, Addgene plasmid # 12260), pCMV-VSV-G (a gift from Robert Weinberg, Addgene plasmid # 8454) were transfected into HEK293T cells using lipofectamine 3000 reagent according manufacturer’s protocol. 48h after transfection, viral supernatants were collected, filtered, mixed (except SCB), and applied to Y1 cells after addition of 8 μg/ml of polybrene (Santa Cruz Biotechnology sc-134220). 48h after Y1 transduction, cells were selected with 3 μg/ml of puromycin for 7 days before testing knock out efficiency.

### 2.9 Clonogenic and viability assays

The indicated amounts of cells per well were plated on 6-wells or p60 plates (figure 4B only), let to adhere overnight and then treated as described. After that, the culture media were replaced every other day until the end point. Cells were then washed with PBS, fixed and stained in a fixing/staining solution (0.05% crystal violet, 1% formaldehyde, 1% methanol in PBS) and washed abundantly. Images were acquired using GelCount colony analyser (Oxford Optronics, Oxford, UK).

### 2.10 Growth curves

At day 0, 3×10^4^ cells per 35 mm culture dish were plated in DMEM-FCS medium with or without FGF2. At the indicated days, cells were harvested in triplicates, fixed in formaldehyde 3.7%, diluted in Phosphate Buffered Solution (PBS) and stored. The medium of reminiscent plates was changed in every 2 days. Cells were later counted in a Z2 Beckman Coulter^®^ counter.

### 2.11 Non-adherent proliferation assay

At day 0, 1×10^4^ cells per well were plated on ultra-low attachment 96 wells plates (Corning CLS3474). Relative cell viability/proliferation was measured after 1 day, to set up a baseline, and after 10 days using CellTiter 96 AQ_ueous_ (Promega G3582) according to the manufacturer’s protocol. A least 9 wells per cell were assayed at each time point.

### 2.12 Statistical analyses

Bar graphs with two columns were analyzed with paired Student’s t-test and bar graphs with tree or more columns were analyzed by one-way ANOVA of variance followed by multiple comparison post-test. Growth curves were analyzed by two-way ANOVA of variance followed by multiple comparison post-test. All graphics and statistical analyses were performed using GraphPad Prism 7 software.

## 3 RESULTS

### 3.1 FGF2 impairs cell cycle progression in K-Ras-driven cancer cells

We previously showed that FGF2 triggers G0/G1→S transition but irreversibly restrains the proliferation of K-Ras-driven Y1 malignant cells (Costa et al., 2008). Y1 cells are poorly synchronized by serum starvation. Hence, to address FGF2 effects along cell cycle progression accurately, we initially used the Y1 D1 subline. These cells, which we described elsewhere (Schwindt et al., 2003), display strict control of quiescence/proliferation switch in response to serum, and are phenotypically identical to parental Y1 cells regarding karyotype, K-Ras overexpression and malignant phenotype.

Initially, we followed cell cycle progression of G0/G1 arrested Y1D1 cells after serum stimulation +/- FGF2, collecting samples every 2h, with a 30min pulse of BrdU uptake into DNA, immediately before cell harvesting (Figure 1A and 1B). Flow cytometry results showed that FGF2 delayed both cell entry in and progression through S phase (Figure 1B, upper and middle panel). In FGF2-treated samples, after 20h of stimulation, we observed BrdU unlabeled S-phase cells, indicating DNA synthesis arrest (Figure 1A arrows). Moreover, between 24 and 48h, we found a parallel decrease in S-phase and accumulation in G2/M sub-population (Figure 1B, middle and lower panel). This accumulation likely was due to G2 arrest, since mitosis blockage by colchicine induced G2 and M-phase accumulation in serum-stimulated but not in FGF-2-treated cells between 24 and 36h (Figure 1C, top and middle panel). Notably, about 40% of the FGF2-stimulated cells remained in G0/G1 phase irrespective of colchicine addition (Figure 1B and 1C, bottom panel); indicating that many of the FGF2-stimulated cells were not even able to leave G1 phase. These results showed that FGF2 compromises cell fitness, impairing the progression throughout the cell cycle in these Ras-driven malignant cells.

**Figure 1.**
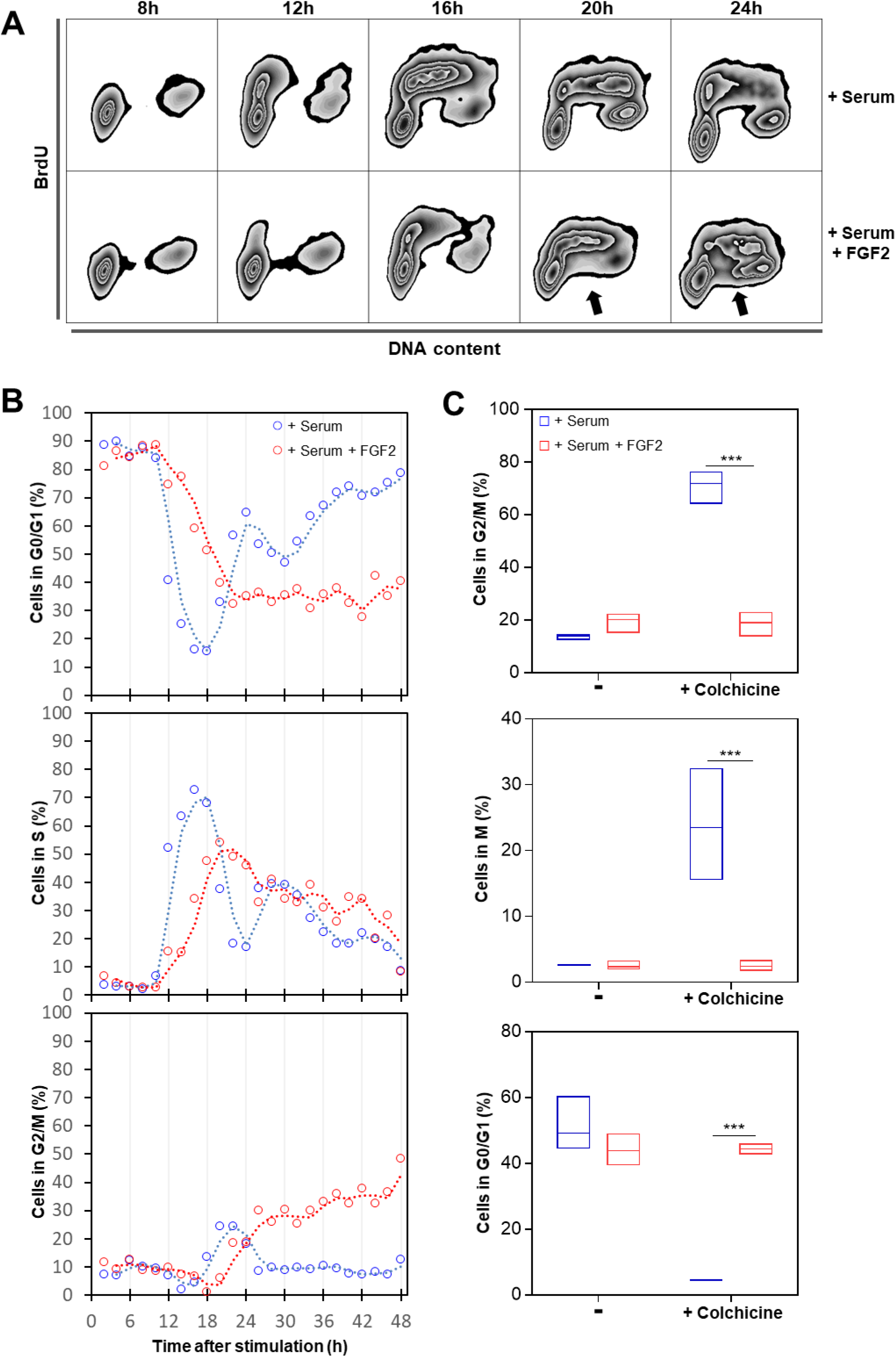
FGF2 impairs cell cycle progression in K-Ras-driven cancer cells. Serum-starved Y1D1 cells were stimulated by 10% serum with or without 10 ng/ml FGF2 to re-entry the cell cycle. Cells were subjected to a BrdU pulse 30 minutes before sample collection (every two hours). **(A)** Representative zebra plot flow cytometry data of BrdU-stained cells versus DNA content at the indicated times after stimulation comparing cell cycle re-entry and progression with or without FGF2. BrdU was added at 50 μM for 30 minutes before harvesting. The arrows indicate BrdU unlabeled S-phase cells. **(B)** Time-course flow cytometry analyses comparing the progression along cell cycle phases from 2h to 48h after stimulation by serum with or without FGF2. BrdU was added at 50 μM for 30 minutes before harvesting. **(C)** Quantifications of flow cytometry data showing phospho-histone H3 (S10) and DNA content double stain. The proportions of cells in each phase were measured 24, 28, 32, and 36h after stimulation with serum or serum + FGF2 in presence or absence of 2μM colchicine. Mitotic cells were addressed by chromatin condensation indicated by phospho-histone H3 (S10) positive stain. Asterisks indicate significant differences. (***) means p ≤ 0.001.

### 3.2 FGF2 exacerbates replication stress and sensitizes K-Ras-driven cancer cells to checkpoint inhibition toxicity

The S-phase cells displaying DNA synthesis arrest in the flow cytometry data (Figure 1A arrows) suggested that FGF2 induced replication stress in this K-Ras-driven cell model. As unresolved replication arrest generates double-strand breaks, we measured the levels of the DNA damage marker phospho-H2AX histone (γ-H2AX) (Gagou et al., 2010) in Y1 cells stimulated by serum +/- FGF2. Serum-stimulated cells exhibited moderate levels of γ-H2AX, and these levels increased over 3.5-fold by FGF2 stimulation (Figure 2A). As a more specific readout of replication stress, we incorporated the thymidine analogue BrdU to these cells and measured single-stranded DNA (ssDNA) foci at native conditions. FGF2 stimulation resulted in about 45% of the cells showing more than 10 ssDNA foci after 24h, comparing to less than 3% of the control cells (Figure 2B and 2C). To further confirm that such DNA damage results from replication stress, we analyzed the distribution of γ-H2AX positive cells along the cell cycle at three different time points after stimulation. Corroborating the above results, serum-stimulated cells displayed moderate γ-H2AX staining, and FGF2 increased γ-H2AX-positive cell population 18 and 24h after stimulation (Figure 2D). In all samples γ-H2AX-positive cells were almost exclusively found in S and G2 phases, pointing that the observed DNA damage in these cells is likely a consequence of replication stress.

**Figure 2.**
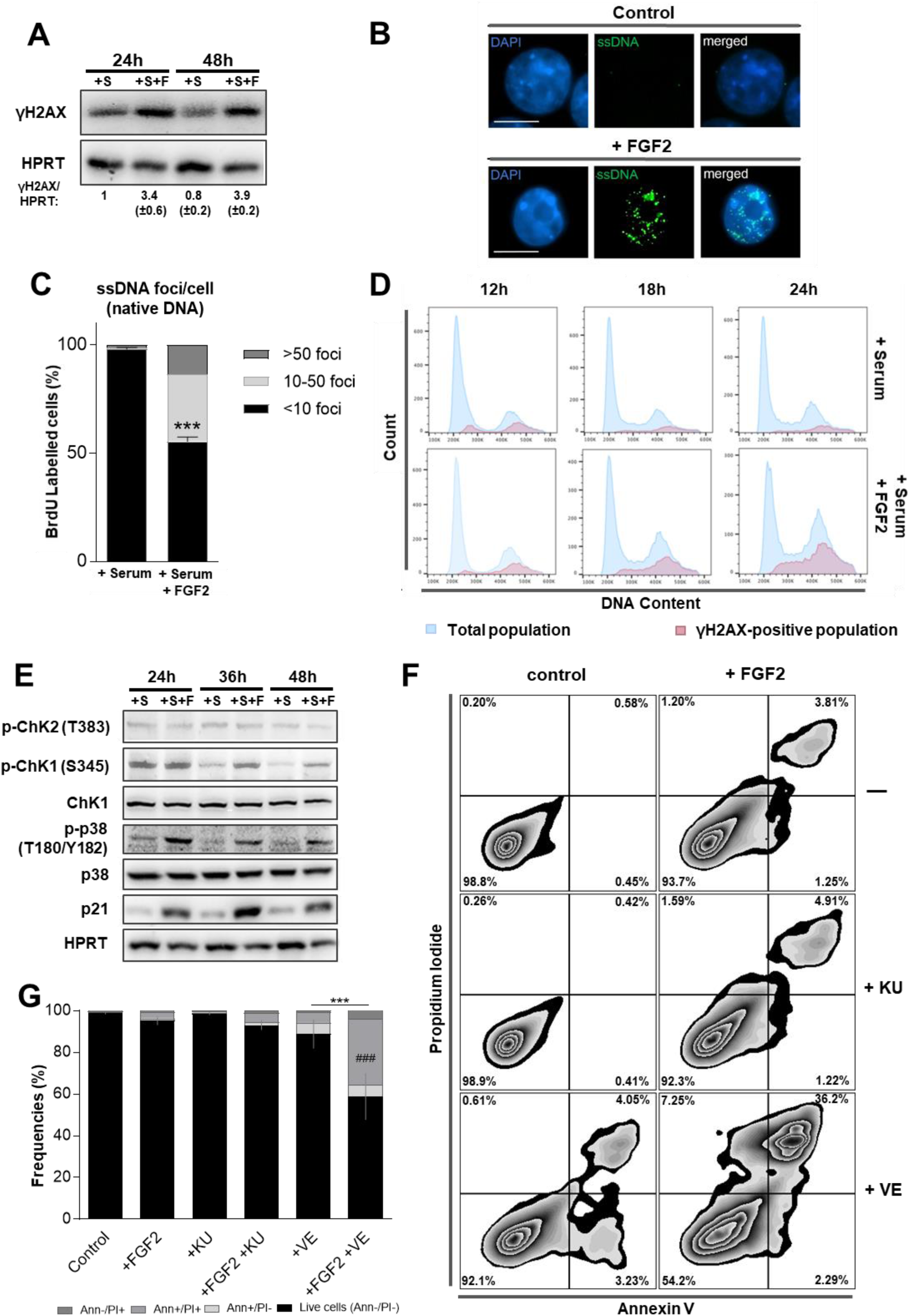
FGF2 reinforces replication stress in K-Ras-driven cancer cells increasing ATR-checkpoint inhibition toxicity. **A)** Western blots comparing the levels of phospho-H2AX histone (γ-H2AX) in Y1 cells. Cells were serum starved and then re-stimulated with 10% serum (+S) or 10% serum + 10ng/ml FGF2 (+S+F) for the indicated times. HPRT was used as a loading control. Quantifications were performed using Uvitec Alliance 9.7 software. **B)** Representative immunofluorescence detection of single-stranded DNA (ssDNA) foci under native conditions. 50mM of BrdU was incorporated to Y1 cells for 48h and then washed out. Cells were grown for additional 24h in complete media with or without 10ng/ml FGF2, and then stained for BrdU (green) and DNA (blue) under non-denaturing conditions. White bars correspond to 10 μm. **C)** Quantification of ssDNA foci per cell from the experiments described in B. Error bars indicate mean ± S.D. This assay was performed in triplicate with measurement of at least 65 cells per replicate (n = 65/assay). Asterisks indicate statistically significant differences. (***) means p ≤ 0.001. **D)** Representative histogram flow cytometry data comparing serum and serum + FGF2 stimulation regarding γ-H2AX distribution along the cell cycle phases. Y1 cells were serum starved and then re-stimulated by 10% serum with or without 10ng/ml FGF2 for the indicated times. **E)** Western blots comparing the levels of the DDR and checkpoint markers phosphorylated Chk1 (p-Chk1), phosphorylated Chk2 (p-Chk2), phosphorylated p38 MAPK (p-p38), and p21 in Y1 cells re-stimulated by 10% serum (+S) or 10% serum + 10ng/ml FGF2 (+S+F) for the indicated times after serum starvation. Total Chk1, p38, and HPRT were used as loading controls. **F)** Representative zebra plot flow cytometry data of Y1 cells growing in complete media in presence or absence of 10ng/ml FGF2 for 48h, with concomitant addition of 5μM of the ATM inhibitor KU-55933 (KU) or the ATR inhibitor VE-821 (VE). Annexin V/propidium iodide (PI) double stain was used to address cell death. **G)** Quantification of the experiments described in F. Error bars indicate mean ± S.D. of live cells (n=3, from independent experiments). Asterisks indicate statistically significant differences. (***) means p ≤ 0.001. (#) refers to significant differences from the FGF2-treated sample.

These observations prompted us to probe for the engagement of the DNA Damage Response (DDR). We reasoned that DDR and checkpoint activation might contribute to the observed cell cycle arrest triggered by FGF2 in these cells, as a protective response against FGF-induced replication stress. We first assessed the activation status of classical DDR and checkpoint effector proteins, i.e., Chk 1 and 2, p21 and p38 (Reviewed by Harper & Elledge, 2007). The results showed that FGF2 does not alter the levels of active Chk2; however, FGF2 increased the levels of phosphorylated Chk1, p38 and p21 comparing to serum-stimulated samples (Figure 2E). These results confirmed that FGF2-induced replication stress triggers DDR and checkpoint activation in these cells.

Checkpoint inhibition prevents cell cycle arrest induced by DNA damaging chemotherapy, forcing cancer cells into a defective mitosis and consequent cell death (Huntoon et al., 2013). To assess whether combined FGF2 signaling and checkpoint inhibition could lead to this same outcome, we focused on ATM and ATR, the two major kinases controlling checkpoint response (Harper & Elledge, 2007). We treated cells with specific ATR (VE-821) (Reaper et al., 2011) and ATM (KU-55933) (Hickson et al., 2004) pharmacological inhibitors for 48h in the presence or absence of FGF2, and measured cell death on the flow cytometer. FGF2 only modestly increased cell death compared to serum-stimulated control samples. ATM inhibition had no significant effect on cell death with or without FGF2. ATR inhibition alone moderately increased cell death. Strikingly, the association of VE-821 and FGF2 induced over 40% of cell death after 48h (Figure 2F and 2G).

Altogether, these results indicate that Y1 cells, as proposed for other cancer models, deal with chronic replication stress and rely on checkpoint activation for cell survival. FGF2 signaling upregulated such basal replication stress leading to cell cycle arrest and, at the same time, increasing checkpoint dependence. Thus, FGF2 sensitizes this K-Ras-driven malignant model to cell death induced by ATR-mediated checkpoint inhibition.

### 3.3 FGF2 induces proteotoxic stress and sensitizes K-Ras-driven cancer cells to bortezomib cytotoxicity

In agreement with our previous report (Costa et al., 2008), flow cytometry data from FGF2-treated cells presented on figure 1 showed increased cell size (FSC) and internal complexity (SSC) comparing to serum-stimulated control cells (Figure 3A). To further investigate this dual effect of FGF2 blocking proliferation but keeping cells growing, we stimulated serum starved Y1 cells with serum +/-FGF2 and measured average size and the amount of protein per cell. FGF2-stimulated cells displayed increased cell size and about twice the amount of protein/cell measured on serum-stimulated samples after 48 and 72h (Figure 3B). These results indicated that although FGF2 triggered cell cycle arrest in Y1 cells, it stimulated cell growth concerning volume and mass.

**Figure 3.**
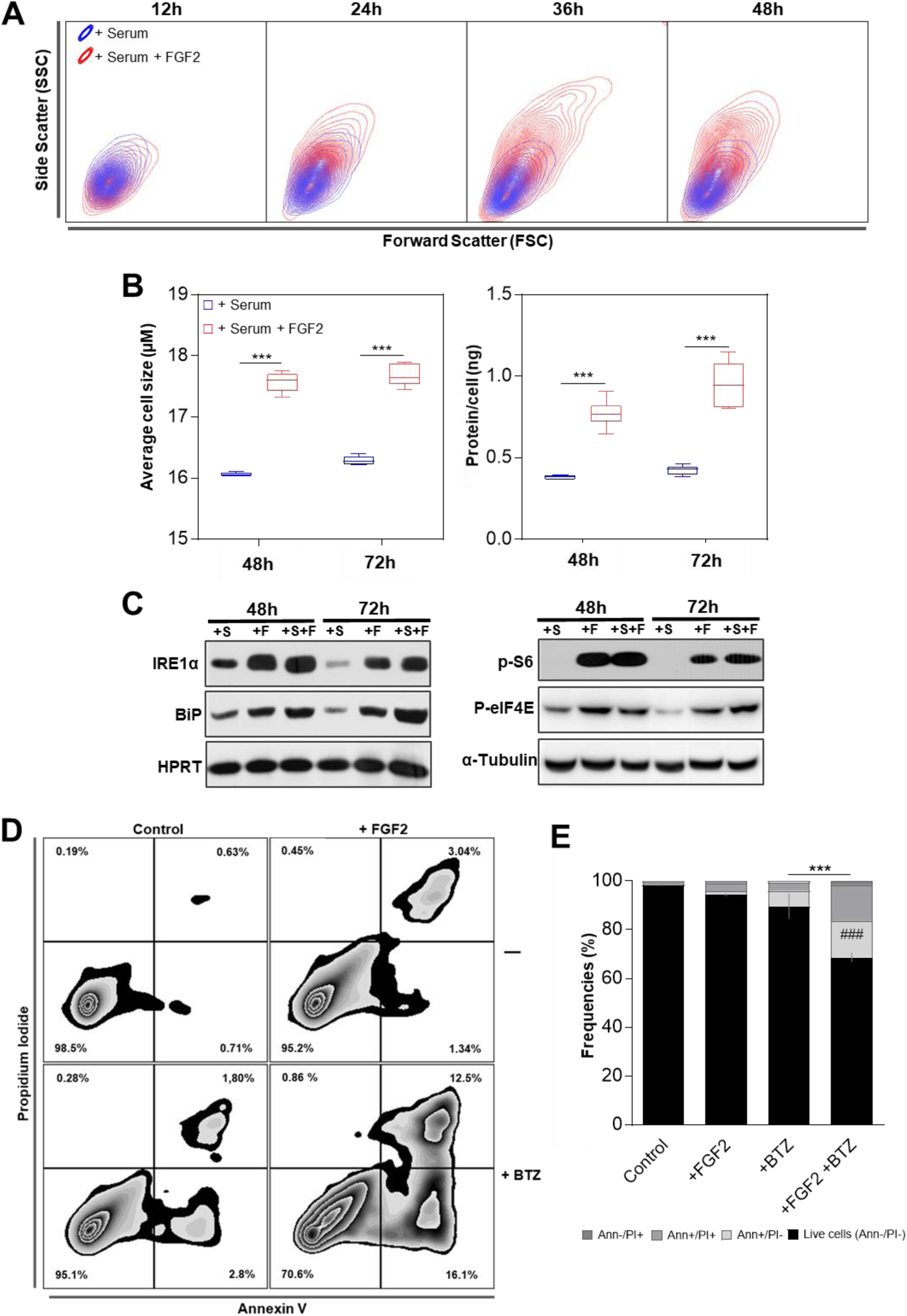
FGF2 disrupts the proteostasis and sensitizes K-Ras-Driven cancer cells to bortezomib toxicity. **(A)** Representative contour plot flow cytometry data from samples of figure 1B comparing serum and serum + FGF2 stimulation regarding cell size (Forward Scatter) and internal complexity (Side Scatter) along the time. **(B)** Measurements of the average cell size and the amount of protein per cell comparing serum and serum + FGF2 stimulated cells. Y1 cells were serum starved and then re-stimulated by the indicated times. Asterisks indicate significant differences. (***) means p ≤ 0.001. (n=6, from independent experiments) **(C)** Western blots comparing the levels of the UPR markers IRE1α and Bip (left panel); and the phosphorylated forms of S6 ribosomal protein (p-S6) and eukaryotic translational initiation factor 4E (p-EIF4E; right panel) among the different stimuli. Y1 cells were serum starved and then re-stimulated with 10% serum (+S); 10ng/ml FGF2 (+F); or both (+S+F) for the indicated times. HPRT and α-tubulin were the used as loading controls. **(D)** Representative zebra plot flow cytometry data of Y1 cells growing in complete media in presence or absence of 10ng/ml FGF2 for 96h, with the addition of 20 nM bortezomib (BTZ) in the last 72h. Annexin V/propidium iodide (PI) double stain was used to address cell death. **(E)** Quantification of the experiments described in **D**. Error bars indicate mean ± S.D. of live cells (n=3, from independent experiments). Asterisks indicate statistically significant differences. (***) means p ≤ 0.001. (#) refers to significant differences from the FGF2-treated sample.

The rates of protein synthesis and degradation show physiological fluctuations; however, an optimal balance between these processes is required to warrant cell viability (Walter & Ron, 2011). We then investigated whether this protein overload induced by FGF2 results in proteotoxic stress and, consequently, Unfolded Protein Response (UPR) activation. To this end, we measured the levels of the endoplasmic reticulum kinase IRE1α and the molecular chaperone Bip, two core sensors of UPR, which are upregulated in cells facing proteotoxic stress (Ron & Walter, 2007). We found increased levels of both proteins at 48 and 72h-FGF2-stimulated cells, even in serum-free media, comparing to serum-stimulated control samples (Figure 3C, left). Interestingly, despite this active UPR, FGF2-stimulated cells displayed high levels of both, phosphorylated S6 ribosomal protein and eukaryotic translational initiation factor 4E (eIF4E) (Figure 3C, right). The phosphorylated forms of these proteins indicate active protein synthesis (Sonenberg & Hinnebusch 2009); implying that FGF2 aggravates the proteotoxic stress by maintaining active protein synthesis irrespective of an ongoing activated UPR.

The enhanced proteotoxic stress of malignant cells is a clinical target in cancer therapy (Csizmar et al., 2016). Hence, we tested if, beyond the cytostatic effect as a single agent, FGF2 could sensitize Y1 cells to the cytotoxicity of proteasome inhibition. We treated the cells with FGF2 for 24h and then added bortezomib (BTZ) for additional 72h before harvesting. We then measured cell death by annexin V/Propidium iodide double stain on the flow cytometer. In the absence of FGF2, Y1 cells were very tolerant to 20nM of BTZ for 72h, showing almost 90% of live cells. In contrast, the combination of FGF2 and BTZ reduced the percentage of live cells to less than 70%, while FGF2 alone had only minor effects on cell death (Figure 3D and 3E). These observations not only show that FGF2 stress response disrupts the proteostasis, but also that it can be combined with proteasome inhibition to trigger cell death in these K-Ras-driven cancer cells.

### 3.4 K-Ras depletion prevents FGF2 toxicity and sensitization to checkpoint or proteasome inhibition in K-Ras-driven cancer cells

The malignant phenotype of Y1 cells is attributed to the overexpression of the wild-type K-Ras protein resulting in high basal levels of K-Ras-GTP (Schwab et al., 1983). Using the isoform unspecific RasN17 dominant negative, we previously proposed that FGF2 toxicity in Y1 cells depends on high basal levels of K-Ras-GTP (Costa et al., 2008). To link causally the high levels of K-Ras protein to FGF2 toxicity and sensitization to stress-targeted inhibitors, we performed Cas9-mediated genome editing to deplete K-Ras protein in these cells. After antibiotic selection, the resultant polyclonal subline (hereafter Y1-3K) displayed more than 10-fold reduction in K-Ras protein levels, comparing to the scramble-transduced control cells (hereafter Y1-scb) or the parental Y1 cells (Figure 4A).

**Figure 4.**
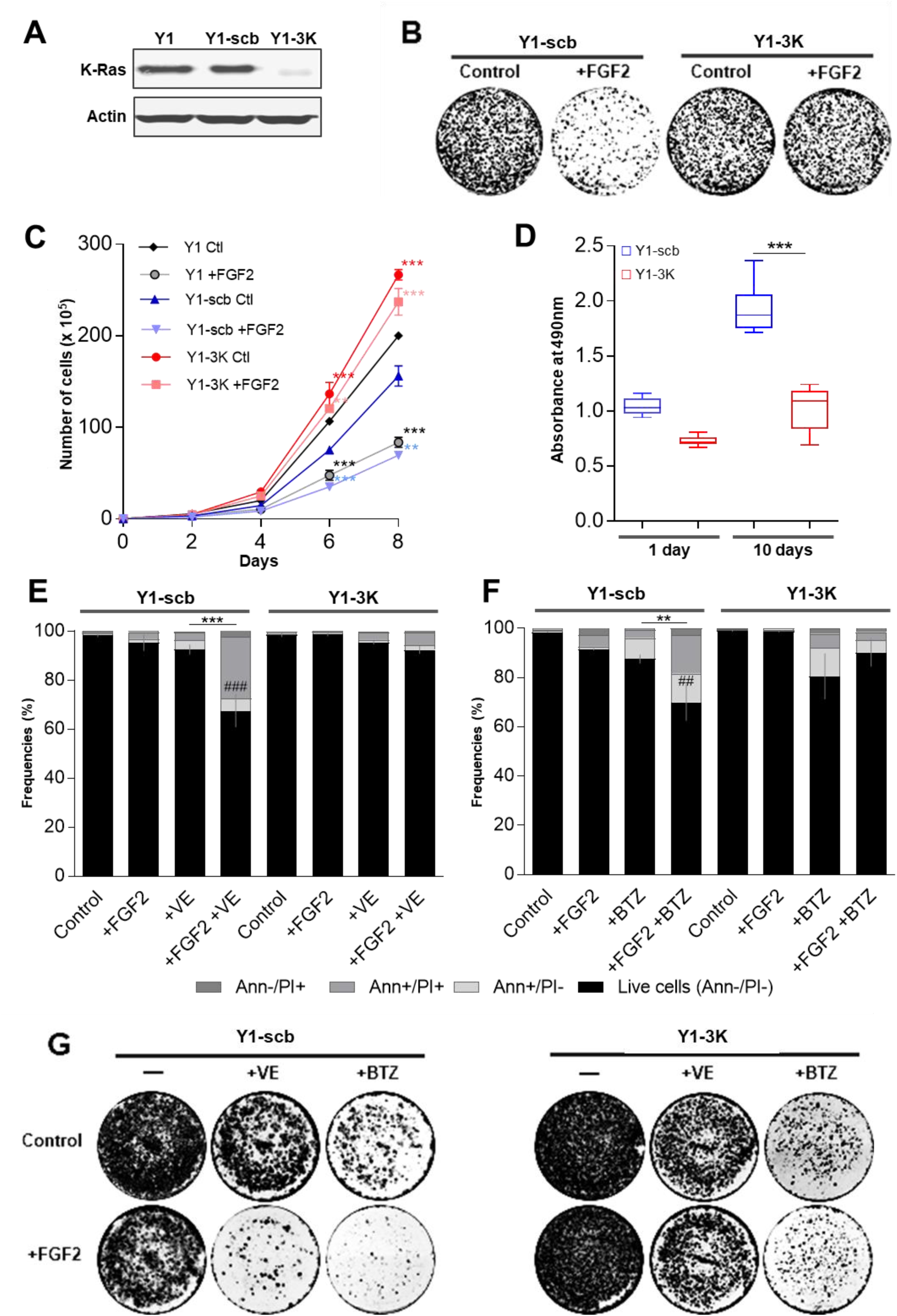
K-Ras overexpression is required for FGF2 toxicity and sensitization to cell death induced by checkpoint or proteasome inhibition. **(A)** Western blots comparing the levels of K-Ras among Y1 parental, Y1-scb control, and Y1–3K K-Ras depleted cells. Lysates were prepared from cells growing at complete media. Actin was used as a loading control. **(B)** Representative clonogenic assays comparing the viability of Y1-scb and Y1–3K cells. For each cell line, 120 cells/cm^2^ were plated in complete media in presence or absence of 10 ng/ml FGF2, grown for 15 days, and then fixed/stained. Culture media and FGF2 were renewed every 2 or 3 days. **(C)** Representative growth curves comparing the proliferation of Y1 parental, Y1-scb control, and Y1–3K K-Ras depleted cells. For each cell line, 3×10^4^ cells were plated in complete media in presence or absence of 10 ng/ml FGF2 and grown for the indicated times. Culture media and FGF2 of the reminiscent plates were renewed at every harvest point. Error bars indicate mean ± S.D. of technical triplicates. Asterisks indicate significant differences from Y1 control condition. (**) means p < 0.01 and (***) means p ≤ 0.001. **(D)** Non-adherent proliferation assay comparing Y1-scb and Y1–3K cells. For each cell line, 1×10^4^ cells were plated on ultra-low attachment 96 wells plates in complete media. Relative cell viability/proliferation was addressed after 1 day, to set up a baseline, and after 10 days to measure proliferation using CellTiter 96 AQ_ueous_ (Promega). At least 10 wells per cell were assayed at each time point. Asterisks indicate significant differences. (***) means p < 0.001. **(E)** Flow cytometry data of Y1-scb control, and Y1–3K K-Ras depleted cells growing in complete media in presence or absence of 10ng/ml FGF2 for 48h, with concomitant addition of 5μM VE-821 (VE). **(F)** Flow cytometry data of Y1-scb control, and Y1–3K K-Ras depleted cells growing in complete media in presence or absence of 10ng/ml FGF2 for 96h, with the addition of 20nM bortezomib (BTZ) in the last 72h. For **E** and **F**, Annexin V/propidium iodide (PI) double stain was used to address cell death. Error bars indicate mean ± S.D. of live cells (n=3, from independent experiments). Asterisks indicate statistically significant differences. (**) means p ≤ 0.01 and (***) p ≤ 0.001. (#) refers to significant differences from the FGF2-treated sample. **(G)** Representative assays comparing the long-term viability of Y1 -scb and Y1–3K cells treated with the combinations of FGF2 and VE-821 or bortezomib. For each cell line, approx. 2,8×10^4^ cells/cm^2^ were plated and treated as described in **E** and **F**. After the treatments, the stimuli were washed out, the plates were grown in complete media for additional 10 days, and then fixed/stained.

We next enquired whether K-Ras depletion impacted on viability and proliferation of Y1 cells, as well as its likely protective effect from FGF2 toxicity. Clonogenic assays showed no significant change in cell viability caused by K-Ras depletion (Figure 4B). Moreover, K-Ras depletion prevented FGF2 toxic effects on long-term viability of Y1-3K cells (Figure 4B). Furthermore, growth curves indicated that Y1-3K cells proliferate significantly faster than both parental Y1 and Y1-scb control cells, and FGF2 did not impact on cell proliferation of this K-Ras depleted subline (Figure 4C). Conversely, K-Ras depletion restrained the proliferation of Y1-3K cells under non-adherent growth conditions (Figure 4D). This set of results shows that K-Ras depletion elicited robust survival and proliferation in solid substrate but suppressed some malignant phenotype traits in this cell model.

We then investigated whether K-Ras depletion is sufficient to prevent the cell death induced by simultaneous FGF2 stimulation and proteasome or ATR-checkpoint inhibition in these cells. To this end, we treated Y1-scb and Y1-3K cells with combinations of FGF2, bortezomib and VE-821 in the same regimens described above, and measure cell death by flow cytometry. The results for Y1-scb cells, as expected, were similar to those shown for Y1 parental cells; with the combinations of FGF2 with VE-821 or bortezomib inducing over 30% of cell death (Figure 4E and 4F, left). On the other hand, in Y1-3K cells, K-Ras depletion largely prevented the cell death induced by the combination of FGF2 and VE-821 (Figure 4E, right). Strikingly, in these K-Ras-depleted cells, bortezomib induced about 25% of cell death, and the presence of FGF2 partially rescued these cells from bortezomib cytotoxicity (Figure 4F, left). To address the effects of these toxicities on long-term cell viability, we treated both cells using the same regimens described above, washed out FGF2 and the inhibitors and cultured the cells for additional 10 days. In agreement with the flow cytometry results, combinations of FGF2 and VE-821 or bortezomib strongly reduced the long-term viability of Y1-scb cells (figure 4G, left); and K-Ras depletion fully prevented these toxicities in Y1-3K cells (figure 4G right). Altogether, these data indicated that K-Ras overexpression underlies FGF2 toxicity and the sensitization to proteasome or ATR checkpoint inhibition in these cells. Moreover, they highlight that FGF2 signaling can be toxic or cytoprotective depending on the context; in this case, levels of K-Ras expression and malignant phenotype.

### 3.5 FGF2 triggers sustained MAPK-ERK1/2 overactivation and lethally sensitizes human cancer cells to proteasome and checkpoint inhibitors

The above data, focused on K-Ras-driven murine Y1 cancer cells, implied that mitogenic signaling activation combined with stress-response pathways inhibitors could disrupt the homeostatic robustness of cancer cells resulting in cell death. Hence, we asked whether this hypothesis would hold true for human cancer cells. Cytostatic and cytotoxic effects of FGF2 over ESFT cells have been reported by different researchers in the last decades, with the specific molecular mechanisms of this toxicity varying among these studies (Schweigerer et al., 1987; Williamson et al., 2004; Passiatore et al., 2011). Thus,we tested A673, RD-ES, SK-N-MC and TC-32 ESFT cells for the toxicities of these combinations of FGF2 and proteasome or checkpoint inhibitors. We focused on the potential of these regimens to kill cancer cells, irrespectively of the cell death subroutine engaged, as well as to reduce the number of viable cancer cells; using concentrations in which none of these stimuli induce pronounced cell death as a single agent. Thus, for each cell line, we plated the indicated number of cells and, after treatments, we measured cell death by annexin V/Propidium iodide double stain and the total number of cells in each sample by flow cytometry. For proteasome inhibition, we treated the cells with FGF2 for 24h and then added bortezomib for additional 48h before harvesting. The regimen for checkpoint inhibition was 72h treatment with FGF2 combined with ATM (KU-55933), ATR (VE-821) or both inhibitors; since the functions of these kinases can overlap but are not redundant (Cimprich & Cortez, 2008). Overall, the combination of FGF2 and proteasome or checkpoint inhibition increased the cell death and reduced the number of live cells to a greater extent than the respective stimuli as single agents in all four cancer cell models (figure 5A). For proteasome inhibition, striking results were found in A673, SK-N-MC and TC-32 cells, in which the association with FGF2 reduced the number of live cells to about 1/3 of the observed for bortezomib alone (figure 5A). In all ESFT cells, the association of FGF2 and VE-821 reduced the number of live cells to about half of the measured using this ATR inhibitor as a single agent (figure 5A). While the association of FGF2 and KU-55933 resulted in significant increased toxicity only in A673 cells (figure 5A); in agreement with the role of ATR as the major player in the replication stress response (Saldivar et al, 2017).

**Figure 5.**
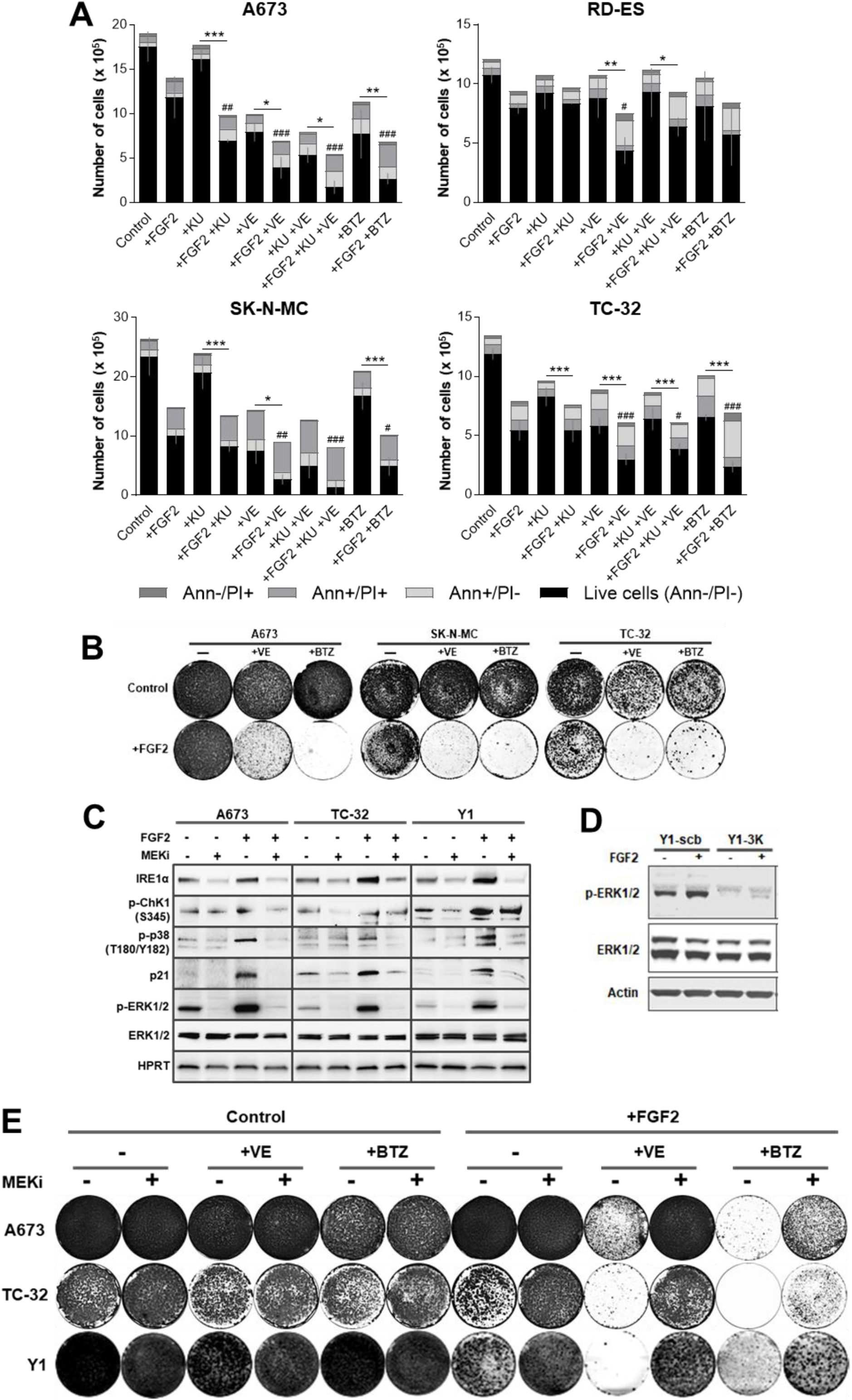
FGF2 promotes MAPK-ERK1/2 sustained overactivation and lethally sensitizes human cancer cells to checkpoint and proteasome inhibition. **(A)** Flow cytometry data of ESFT cells growing in complete media in presence or absence of FGF2 for 72h, with concomitant addition of KU-55933 (+KU), VE-821 (+VE), both (+KU +VE), or addition of bortezomib (+BTZ) in the last 48h. Concentrations as following: A673 cells FGF2 20ng/ml, KU 5μM, VE 5μM, and BTZ 10nM; 1×10^5^ cells were plated. RD-ES cells FGF2 20ng/ml, KU 5μM, VE 5μM, and BTZ 10nM; 2,5×10^5^ cells were plated. SK-N-MC cells FGF2 1ng/ml, KU 5μM, VE 2μM, and BTZ 10nM; 2.5×10^5^ cells were plated. TC-32 cells FGF2 5ng/ml, KU 5μM, VE 2μM, and BTZ 10nM; 1,5×10^5^ cells were plated. Annexin V/propidium iodide (PI) double stain was used to address cell death. Results are expressed in absolute numbers of cells per plate 72h after stimulation. Error bars indicate mean ± S.D. of live cells (n=3, from independent experiments). Asterisks indicate statistically significant differences. (*) means p < 0.05, (**) p < 0.01 and (***) p ≤ 0.001. (#) refers to significant differences from FGF2-treated sample. **(B)** Representative assays comparing the long-term viability of A673, TC-32 and SK-N-MC cells treated with the combinations of FGF2 and VE-821 or bortezomib. Cells were plated and treated as described in **A**. After the treatments, the stimuli were washed out, the plates were grown in complete media for additional 10 days, and then fixed/stained. **(C)** Western blots comparing the levels of phospho-ERK1/2 (p-ERK1/2) and the stress markers IRE1α, phosphorylated Chk1 (p-Chk1), phosphorylated p38 MAPK (p-p38), and p21 among A673, TC-32 and Y1 cells in the presence or absence of FGF2 and the MEK inhibitor selumetinib. FGF2 (+) (20 ng/ml for A673; 5 ng/ml for TC-32 and 10 ng/ml for Y1 cells) was added to cells growing at complete media and 5 μM selumetinib was added to the indicated plates (MEKi +) 8h after FGF2 addition. Plates were harvested 24h after FGF2 addition. Total ERK and HPRT were used as loading controls. **(D)** Western blots comparing the levels of phospho-ERK1/2 (p-ERK1/2) among Y1-scb and Y1-3K cells in the presence or absence of FGF2. FGF2 (+) (10 ng/ml) was added to cells growing at complete media and plates were harvested 24h later. Total ERK and actin were used as loading controls. **(E)** Representative assays comparing the long-term viability of A673, TC-32, and Y1 cells treated with the combinations of FGF2 and VE-821 or bortezomib, with or without the addition of the MEK inhibitor selumetinib. Cells were plated and treated as described in **A**. 8h after FGF2 stimulation, 5μΜ of selumetinib (MEKi +) was added to the indicated plates. 72h after FGF2 addition, the stimuli were washed out, the plates were grown in complete media for additional 10 days, and then fixed/stained.

We next addressed the effects of proteasome or ATR-checkpoint inhibition on long-term cell viability with or without FGF2. We treated A673, SK-N-MC and TC-32 cells as described above, washed out FGF2 and the inhibitors and cultured the cells for additional 10 days. The results show minor or no effects of FGF2, bortezomib or VE-821 after these time for all three cells. Conversely, the associations of FGF2 with these inhibitors were even more toxic than anticipated by the flow cytometry data, resulting in a massive reduction in cell viability after 10 days (figure 5B). These data demonstrated that FGF2 signaling activation can also sensitize human cancer cells to proteasome or checkpoint therapeutic inhibitors.

Ras, EWS-FLI-1 and many other driver oncogenes rely on aberrant MAPK-ERK signaling pathway activation to promote tumorigenesis (Silvany et al., 2000; Dhillon et al., 2007). Thus, we argued whether further overactivation of this same pathway might underlie FGF2 toxicity and the observed increased mobilization and dependence on stress pathways. Our results showed that FGF2 signaling sustains higher levels of p-ERK1/2 even 24h after stimulation, comparing to control cells grown in complete media, in A673, TC-32, and Y1 cells. (figure 5C). This sustained MAPK-ERK activation correlates with the upregulation of UPR (IRE1α) and DDR (p-Chk1, p-p38 and p21) markers (figure 5C). Pharmacological inhibition of MEK 1/2 even 8h after FGF2 stimulation turned off sustained ERK activation and restored cell homeostasis (figure 5C). Coherently, K-Ras depletion, which we showed above to protect from FGF2 toxicity and sensitization to proteasome and checkpoint inhibition, also prevented sustained MAPK overactivation in Y1-3K cells (figure 5D). Finally, we used the same regimens described for figure 5B and added theMEK1/2 inhibitor 8h after FGF2 stimulation. Disruption of FGF2-induced sustained MAPK signaling alleviated or prevented the long-term toxicity triggered by the combinations of FGF2 and ATR-checkpoint or proteasome inhibition in A673, TC-32, and Y1 cells. (figure 5E). Altogether, these results show that, by sustaining the overactivation of MAPK-ERK1/2, a signaling pathway frequently overridden by the malignant transformation, FGF2 reinforces the dependence on stress response pathways, increasing the toxicity of stress-targeted therapeutic inhibitors in both murine and human cancer cells.

## 4 DISCUSSION

Identification and effective targeting of stresses inherent to the malignant phenotype is a current goal for cancer research and therapy. The core rationale of this approach is that uncontrolled malignant proliferation comes with a cost: a stressed phenotype comprising a risky balance between antagonistic metabolic and molecular signaling pathways controlling homeostasis and viability. Therefore, both, further induction and inhibition of stress response pathways can push cancer cells over an irreversible lethal threshold while sparing the normal cell counterparts. In this context, the results presented here highlight how mitogenic signaling can be manipulated to overload inherent stresses, disrupting the risky homeostatic robustness of cancer cells and sensitizing them to stress-targeted therapies.

Many different mechanisms by which growth factors’ mitogenic signaling pathways contribute to the malignant progression have been emphasized in the cancer literature, and gain-of-function mutations along these mitogenic pathways are recognized driver oncogenic lesions in most human cancers. On the other hand, evidence is also accumulating showing that increasing mitogenic activation is not necessarily better for cancer cell fitness. For instance, EGFR and KRAS genes are frequently mutated in lung adenocarcinomas but with no overlap in individual samples. Unni and co-workers have recently shown that synthetic lethality rather than redundancy underlies this mutual exclusivity (Unni et al., 2015). In melanomas, some BRAF V600E-driven tumors become resistant to the BRAF inhibitor vemurafenib through overexpression of BRAF V600E. In this context, withdrawal of the inhibitor resulted in tumor regression caused by a now over activated MAPK pathway (Thakur et al., 2013). These observations suggest that not only the inhibition but also the over-activation of canonically mitogenic pathways might be considered to disrupt tumor cell fitness.

As for other growth factors, pro-tumor roles have been attributed to Fibroblast Growth Factor 2 signaling in different models and contexts (Reviewed by Turner & Grose, 2010). However, most of these results are based on established cancer models, in which an optimal FGF2 signaling level was selected during malignant progression and is now part of its adapted and robust phenotype. Conversely, exogenous administration of FGF2 induced cytostatic or cytotoxic effects in breast cancer (Wang et al., 1998), Ewing’s sarcoma family tumor (Williamson et al., 2004), and medulloblastoma (Fogarty et al., 2007) cell models among others. In vivo, transgenic mice overexpressing FGF2 in all major organs developed into old age showing no increased tumorigenesis (Coffin et al., 1995). Moreover, regular subcutaneous injections of FGF2 also decreased or prevented xenograft tumor growth in mice without noticeable toxicity (Sturla et al., 2000; Costa et al., 2008). These observations suggest that, while FGF2 signaling can be pathologically overridden in certain cancers, exogenous FGF2 administration can disrupt cancer cell homeostasis both in vivo and in vitro.

Accordingly, our time course analyses here showed that FGF2 induces a general, rather than phase-specific, cell cycle arrest in Y1 K-Ras-driven cancer cells. The observed delayed S-phase entry and progression, G1 and G2 arrests likely contribute incrementally to FGF2 cytostatic effects on these cells. This cell cycle arrest results in increased average cell size and protein concentration; implicating that in this model, FGF2 signaling “uncouples” cell growth from proliferation. It is known that EIF4E and S6K signaling play key roles in active protein synthesis and cell size control (Fingar et al., 2002). Thus, the resultant UPR activation is likely a consequence of the sustained EIF4E and S6K activity observed in FGF2-treated cells. One of the strategies of UPR to mitigate proteotoxic stress is to downregulate protein synthesis, helping to restore protein homeostasis (Ron & Walter, 2007). By eliciting active protein synthesis during an ongoing UPR, FGF2 might push proteotoxic stress over the viability threshold. Proteotoxic stress is recognized as a potential Achilles’ heel of malignant cells. Strikingly, treatment of multiple myeloma with bortezomib may result in a complete response. This high sensitivity can be attributed to the extensive production of immunoglobulins by multiple myeloma cells, which accumulates due to bortezomib proteasome inhibition leading to a fatal proteotoxic stress (Obeng et al., 2006; Meister et al., 2007). This scenario provides a rationale for the observed induction of cell death triggered by the combination of FGF2 and bortezomib in these murine cancer cells. Noteworthily, FGF2 can also sensitize ESFT cells to bortezomib cytotoxicity. This panel of cancer cells was largely tolerant to 10nM of bortezomib for 48h. However, the results for A673, SK-N-MC, and TC-32 cells, in which FGF2 or bortezomib alone had minor effects on long-term cell viability but their association was highly toxic, highlight the therapeutic potential of this combination for inducing cancer cell death. It is promising because bortezomib can be very toxic to normal cells, limiting its therapeutic window (Chen et al., 2011). Additionally, in K-Ras-depleted Y1-3K cells, FGF2 alleviated bortezomib toxicity, linking the sensitizing effect of FGF2 to the malignant phenotype and suggesting additional benefits of this combination through the pro-survival effects of FGF2 in normal cells. Our results implicate that FGF2 signaling activation can efficiently disrupt proteostasis, resulting in a common vulnerability in cancer cells with diverse origins and driver oncogenic lesions.

The risky balance between oncogenic activity and increased mobilization of the DDR, frequently found in cancer cells, represents the other vulnerability which we explored here to target these malignant cell models. FGF2 induced stalling or collapse of DNA replication forks in Y1 cells along S-phase. Similar replication stress has been shown to occur early in tumorigenesis; when the oncogenic activity causes increased firing of DNA replication origins leading to unscheduled S-phase progression (Hills & Diffley, 2014). In this scenario, consequent DDR activation upregulating checkpoint proteins is an anti-cancer barrier that must be overcome in early malignant transformation (Bartkova et al., 2005; Gorgoulis et al., 2005). Y1 cells, like other malignant cells, displayed tonic levels of DDR activation which are compatible with high proliferation rates. It is noteworthy that FGF2 stimulation increased the activation of checkpoint proteins, reactivating this anti-cancer barrier and restraining cell proliferation in this K-Ras-driven model. By enhancing replication stress on these cells, FGF2 also increased their dependence on checkpoint activity for survival; hence, the combination of FGF2 stimulation and checkpoint abrogation triggered cell death. Importantly, we showed that the same approach is also effective to trigger cell death in the panel of ESFT cells. The combination of FGF2 and VE-821, but not these agents alone, strongly reduced long-term viability of A673, SK-N-MC, and TC-32 cells. The increased dependency on DDR has been described in other cancer cells as an example of non-oncogene addiction (Luo et al., 2009). In this regard, checkpoint inhibition has recently been shown to sensitize cancer cells to radiotherapy and chemotherapy based on DNA damaging agents (Huntoon et al., 2013; Prevo et al., 2012). However, both radiation and genotoxic agents are frequently very harmful also to non-malignant cells. We provided here evidence that mitogenic signaling activation and checkpoint inhibition might represent an efficient combination stress overloading/sensitization to exploit non-oncogene addiction in cancer therapies.

The specific molecular mechanisms of FGF2 toxicity and sensitization to stress-targeted inhibitors likely vary among Y1 and ESFT cells, and engage different cell death subroutines as observed in our data. However, sustained overactivation of the Ras-MAPKs-ERK1/2 axis by FGF2 is a common feature of the vulnerabilities which we emphasized here. K-Ras depletion in Y1-3K cells prevented MAPK-ERK1/2 overactivation induced by FGF2. These cells showed increased proliferation rates in solid substrate and no FGF2 toxicity or sensitization to proteasome or checkpoint inhibition, but K-Ras depletion also suppressed malignant traits of these cells. These data indicate that the tuning of K-Ras-MAPKs activation, which underlies the proliferation and malignancy in these cells, likely is also the molecular target of FGF2 toxicity. In ESFT cells, EWS/FLI-1 fusion protein suppresses Sprouty 1 expression, a negative-feedback regulator of Ras-MAPKs signaling downstream of FGF receptors; and this is proposed to render unrestrained FGF2-induced proliferation in these cells in vitro and in vivo (Cidre-Aranaz et al., 2017). Indeed, constitutive activation of MAPK-ERK1/2 was found in several ESFT cells, and a Ras dominant negative or MAPK-ERK1/2 pharmacological inhibition restrained the transforming activity of EWS/FLI-1 in immortalized fibroblasts (Silvany et al., 2000). Interestingly, FGF2 itself induces EWS/FLI-1 expression in ESFT cells (Girnita et al., 2000). Taken together, these data suggest that, at optimal growth conditions, exogenous FGF2 likely induce a positive feedback loop resulting in sustained and toxic MAPK-ERK1/2 overactivation in these cells. This scenario is supported by our data showing not only that FGF2 induced sustained higher levels of active ERK1/2, but also that MAPK inhibition even 8h after FGF2 stimulus, restored cell homeostasis and rescued ESFT and Y1 cells from the synergic toxicities which we described above.

The data and the background discussed here argue the question of whether, contra-intuitively, growth factors signaling activation might be clinically explored in cancer therapies. Whilst this major question cannot be exhausted in the scope of this current work, the data provided here show that FGF2 can efficiently disturb the homeostasis of cancer cells from different origin and phenotypes, increasing the toxicity of checkpoint and proteasome inhibitors. Importantly, because we focused here on the sensitizing effect of FGF2, we used doses and times in which neither FGF2 nor the inhibitors trigger massive cell death as a single agent. This implies that the overall toxicity of these combinations over cancer cells can be further improved by tailoring the regimens.

### 4.1 Conclusions

Our data provide evidence that additional stimulation of the same signaling pathways overridden by the malignant transformation might further increase the mobilization and dependence on stress response pathways in cancer cells; hence, improving the efficacy and selectivity of stress-targeted therapies. This approach might be particularly useful at relapsed tumors resulting from acquired resistance to MAPK-ERK1/2 inhibitors, but also provides a potential game-changing novel therapeutic perspective for other human cancers.

## General

We thank Professor Susan A. Burchill for providing the ESFT cells; Dr. Shankar Varadarajan’s lab for providing the Annexin V; and Dr. Nicholas Harper for valuable reagents.

## Author contributions

M.H.D., C.S.F. and H.A.A. conceived the rationale of the experimental design and this manuscript, with fundamental insights from M.S.R. and V.N.

M.H.D., C.S.F., L.L.A., M.S.S. and E.C.L. carried out the experiments.

J.D.Z. performed the statistical analyses.

M.H.D. wrote the manuscript with essential contribution from C.S.F. and J.D.Z.; I.A.P and H.A.A guided and edited the manuscript writing; all authors read and approved the manuscript final version.

I.A.P and H.A.A supervised the project.

